# Sound Frequency Predicts the Bodily Location of Auditory-Induced Tactile Sensations in Synesthetic and Ordinary Perception

**DOI:** 10.1101/2022.06.06.495023

**Authors:** Lua Koenig, Tony Ro

## Abstract

For individuals who experience sound-touch synesthesia, sounds consistently evoke strong, localized sensations on the body. Interestingly, visual and auditory information can also evoke bodily sensations in ordinary perception, such as the sight or sound of a mosquito producing phantom sensations of touch. We systematically investigated the functional relationship between sound frequency and several characteristics of induced tactile experiences in synesthetes and controls. Sound frequency strongly predicted the location of tactile sensations on the body in both synesthetes and controls. Synesthetes felt sound-induced touch more frequently and in significantly more concentrated locations on their bodies compared to sound-induced touch perceptions in controls. This spatial distribution of touch reflects a behavioral mapping between tonotopy and somatotopy that suggests the involvement of early, tonotopically- and somatotopically-organized brain areas. Sound frequency also predicted the hardness, roughness, and sharpness of tactile sensations in both groups. Together, these findings highlight a strong similarity between auditory-tactile mappings in synesthetic and ordinary perception, suggesting that synesthesia only differs in the strength of the mappings and therefore is on a spectrum with ordinary perception. Furthermore, these findings offer insights into the possible underlying neural mechanisms of sound-touch mappings, suggesting that they rely on cross-modal neural pathways already in use in ordinary perception.

## Introduction

The subjective experience of touch can arise even in the absence of physical stimulation. For example, we often feel the phantom sensation of a phone buzz in our pockets. In some disorders, such tactile illusions can arise consistently: in phantom-limb pain, for instance, patients continue to feel strong tactile percepts in an amputated limb^1–3^. Tactile illusions are evidence that the neural mechanisms governing the subjective experience of touch can operate independently from those that process physical tactile signals.

An intriguing phenomenon is the occurrence of tactile sensations in response to stimulation in another modality. For instance, the uncomfortable prickling sensation that arises when nails scrape a blackboard illustrates how auditory information can produce perceptions of touch. In synesthesia, the stimulation of one modality not only causes a normal conscious experience, but also induces a perception in another modality. In sound-touch synesthesia, sounds consistently induce evocative, localized sensations on the body. Unlike other forms of synesthesia, this type may rely on the strong pre-existing correspondences between sound and touch in ordinary perception. One sound-touch synesthete recounted that every time she listened to a specific song, it “filled [her] hands with broken glass in a slimy dishcloth.” The neural mechanisms underlying synesthetic tactile sensations are unknown, despite their potential to shed light on the nature of subjective touch perception and multisensory interactions.

Correspondences between sound and touch are relatively unsurprising given the strong homology between auditory and tactile perception. Indeed, both modalities are mechanosensory and respond to pressure-based stimuli that require transduction from frequency-based signals to neural signals, suggesting they may share similar neural mechanisms. In normal perception, correspondences between sound and touch are frequently reported: for instance, people often report feeling low-frequency sounds, such as a deep bass sound when listening to music, as sensations on their body. Could it be that synesthetic correspondences between sound and touch rely on the same neural mechanisms underlying the connection between sound and touch in ordinary perception?

The inducing stimulus in synesthesia activates a sensory representation corresponding to the synesthetic percept, likely via cross-modal pathways^4,5^. In a patient who developed sound-touch synesthesia after a thalamic stroke, sound-induced tactile sensations correlated with activations in secondary somatosensory cortex (S2)^6^. This activation overlapped with the activation associated with consciously perceiving a physical tactile stimulus. A follow-up diffusion tensor imaging (DTI) study demonstrated that this patient had developed increased connectivity between auditory and somatosensory cortices^7^. Importantly, these activations and connections, although less robust and dense, were also measured in control subjects. These findings suggest that sound-induced tactile illusions in synesthesia may rely on the activation of somatosensory representations through pathways normally utilized during auditory-tactile integration in ordinary perception.

Furthermore, the auditory cortex processes sounds tonotopically, whereby it represents low to high frequency sounds along its rostro-caudal axis^8–11^, and the somatosensory cortex processes touch somatotopically, whereby it represents adjacent areas of the contralateral body surface on a medial-to-lateral direction along the postcentral gyrus^12–18^. Could it be that the increased connectivity observed in sound-touch synesthesia represents corresponding anatomical mappings between the auditory and somatosensory areas that underlie its characteristic sound-induced tactile illusions? Demonstrating a reliable and systematic correspondence between sound frequency and the location of touch on the body in synesthesia would provide strong evidence for the existence of anatomical links between the tonotopic and somatotopic cortical maps.

However, the only studies of sound-touch synesthesia to date involved patients who suffered thalamic strokes, which are likely to have caused extensive cortical reorganization, making these results difficult to generalize^6,7,19,20^. A characterization of sound-touch synesthesia in congenital synesthesia, in individuals who have experienced it since childhood, is lacking. Therefore, the current study’s hypotheses were twofold: first, we hypothesized that sound-touch synesthesia is an extension of ordinary perception. As such, we predicted that sound-touch correspondences in synesthesia should also exist in ordinary perception, albeit to a lesser extent. Second, we hypothesized that systematic anatomical mappings between the tonotopically organized auditory cortex and the somatotopically organized somatosensory cortex were responsible for sound-touch synesthesia. We therefore predicted that we would measure a systematic correspondence between sound frequency and the location of synesthetic tactile sensations on the body in synesthetes. To test these hypotheses, we recruited one group of participants with congenital sound-touch synesthesia, as well as a control group, and systematically measured the characteristics of their tactile sensations in response to simple tones of different sound frequencies.

## Results

Synesthete and control groups reported any tactile sensations they felt while listening to sounds ranging from low pitch (100 Hz) to high pitch (15,000 Hz), including the location of the sensations on their body, its intensity, hardness, dryness, smoothness, coldness, and dullness (Figure 1A).

**Figure 1:**
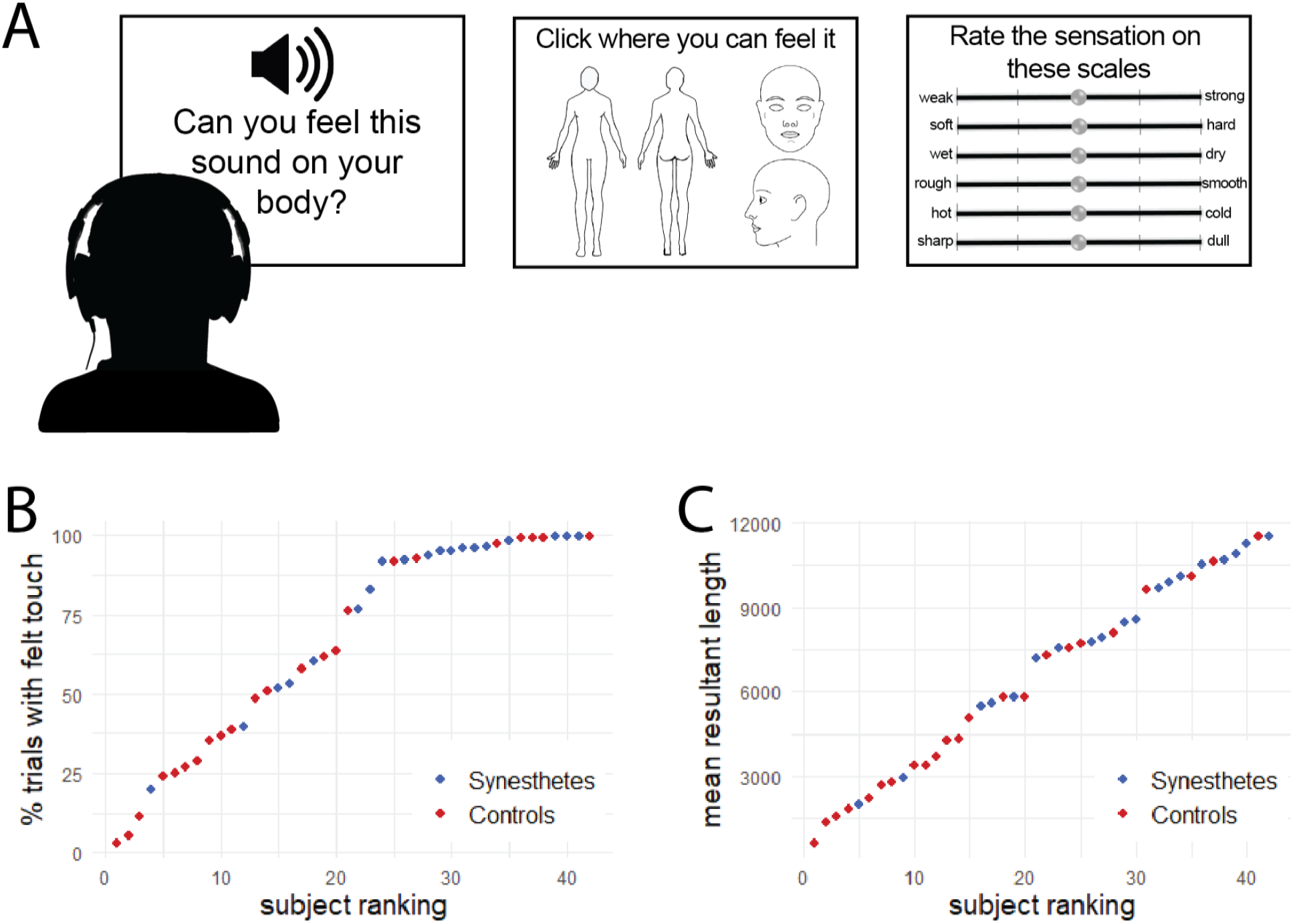
Diagram of the task procedure (A). In each trial, a sound was played repeatedly until the subject indicated whether they felt an accompanying tactile sensation. If so, they indicated where on the body they felt the touch, then rated the sensation on six scales. Synesthetes felt touch in response to sound in a higher percentage of trials compared to controls (B). Overall, synesthetes felt tactile responses to sound more than controls. (C) Synesthetes had higher mean resultant lengths (MRLs) for the location of sound-induced tactile sensations on the body compared to controls, indicating that they felt touch in more concentrated locations on the body.

We first computed the rate of feeling touch in response to sounds in the synesthete and the control groups. To do so, we used a binomial generalized linear mixed-effects model (GLMM) with group and sound frequency as fixed effects, and participant as a random effect. Synesthetes reported feeling touch more frequently (81.1 % of trials, SEM: 1.3%) than controls (55.5% of trials, SEM: 1.5%; main effect of group: *χ*^*2*^*(1)* = 5.83, *p* = 0.016). In both groups, participants reported feeling more sensations of touch as sound frequency increased (main effect of sound frequency: *χ*^*2*^*(12)* = 262.93, *p* < 0.001). Interestingly, at the highest sound frequency, both groups had similar rates of feeling touch (interaction between sound frequency and group: *χ*^*2*^*(12)* = 37.05, *p* < 0.001) (Figure 1B). This dip in the rate of feeling touch for the 15,000 Hz sound likely results from the sound being heard less frequently, as the hearing of high-pitched tones drops in adulthood.

We also assessed how sound frequency influenced the concentration of touch sensations on the body, by examining the polar coordinates of the locations of touch. Specifically, we measured the mean resultant length (MRL) of the selected body locations for each sound frequency. MRL is indicative of the mean concentration of selected locations (see Methods). Larger MRL indicates that the locations of touch are more highly concentrated on the body. Results are presented in Figure 1C and in Table 1. A linear mixed-effects model on the MRL for each sound frequency in each group yielded a main effect of sound frequency (*χ*^*2*^*(12)* = 51, *p* < 0.001) and a main effect of group (*χ*^*2*^*(1)* = 8.85, *p* = 0.0029). A post-hoc correlation between sound frequency and MRL across both groups showed a significant positive correlation (*r*(40) = 0.11, *p* = 0.0138), such that higher frequency sounds elicited less concentrated tactile sensations on the body compared to lower frequency sounds. More importantly for our hypotheses, these results indicate that synesthetes reported sound-induced tactile sensations on more highly concentrated locations of the body.

**Table 1:** Mean resultant lengths (MRL) were computed for each sound frequency and each group. To do so, the vectors of the locations of sound-induced touch selected in each trial were summed and the resulting vector’s length was taken as the MRL larger MRLs correspond to more highly concentrated locations in space, while lower MRLs correspond to more dispersed locations in space.

|  | Synesthetes | Controls |
| --- | --- | --- |
| **100 Hz** | 7052 | 5378 |
| **200 Hz** | 8056 | 4710 |
| **300 Hz** | 8009 | 4644 |
| **400 Hz** | 8086 | 4968 |
| **500 Hz** | 8041 | 5135 |
| **750 Hz** | 7745 | 5780 |
| **1000 Hz** | 8378 | 5172 |
| **1500 Hz** | 7707 | 5761 |
| **2000 Hz** | 8718 | 5685 |
| **3000 Hz** | 8564 | 6328 |
| **5000 Hz** | 9405 | 6445 |
| **10000 Hz** | 8569 | 6567 |
| **15000 Hz** | 7412 | 5399 |

We then evaluated the spatial distribution of sound-evoked touch on the body, in both synesthetes and controls. To do so, we assessed how the location of sound-evoked tactile sensations changed as a function of sound frequency in both groups. We first assessed the correlation between sound frequency and cartesian coordinates along the lateral (X-axis) and the longitudinal (Y-axis) directions on the body (Figure 2). We found no correlation between sound frequency and the location of touch on the lateral body axis in synesthetes (*z*(17) = 0.047, *t* = 0.56, *p* = 0.5853) or in controls (*z*(21) = 0.072, *t* = 0.89, *p* = 0.39). There was no difference in the correlations between sound frequency and the location of touch on the lateral body axis between groups (*t*(40) = -0.22, *p =* 0.83). In contrast, we found a positive correlation between sound frequency and the location of touch on the longitudinal body axis in synesthetes, which was statistically significant (*z*(17) = 0.62, *t* = 5.89, *p* < 0.001). Interestingly, we also found a positive correlation between sound frequency and the locations on the longitudinal axis in controls (*z*(21) = 0.27, *t* = 3.05, *p* = 0.0059). These results indicate that lower frequency sounds were associated with tactile sensations that were significantly lower on the body compared to higher frequency sounds, in both synesthetes and controls. Further, the magnitude of the correlation between sound frequency and the location of touch on the longitudinal body axis was stronger in synesthetes compared to controls (*t*(40) *=* 2.54, *p* = 0.016).

**Figure 2:**
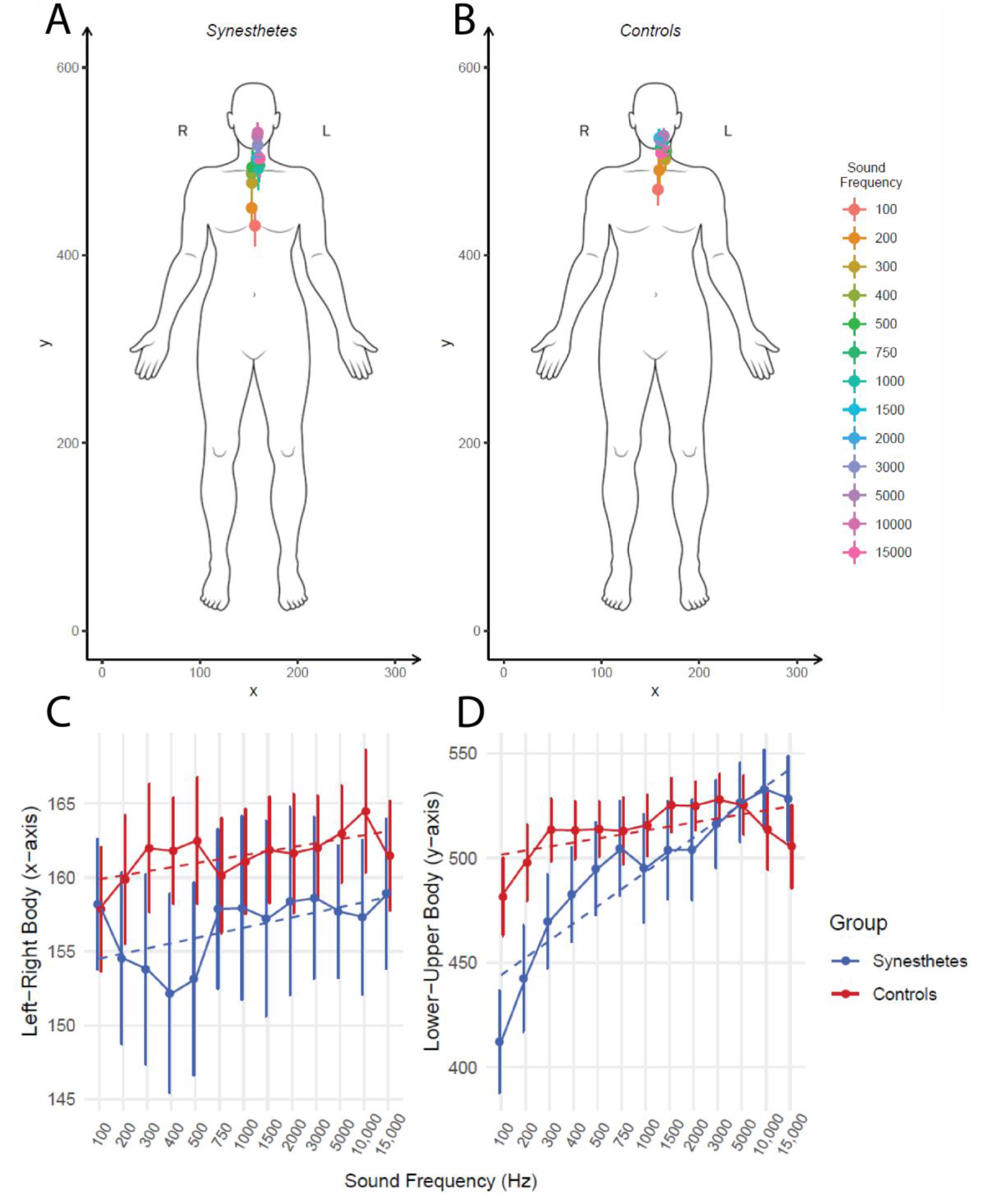
Spatial distribution of sound-induced touch on the body in synesthetes (A) and controls (B). Mean locations on the lateral body axis (C) and the longitudinal body axis (D) are shown across sounds and groups. Dashed lines represent the linear trend across sounds for each group. Error bars represent the standard error of the mean.

Finally, we used linear mixed effects models to assess the effects of sound frequency and group on the qualitative characteristics of sound-induced tactile sensations, with participant and trial number as random effects (see Figure 3). Overall, synesthetes and controls showed similar ratings on the qualitative scales, with no main effect of group, for intensity, hardness, dryness, smoothness, coldness, and dullness. Specifically, the linear mixed-effects model on the intensity of sound-induced touch yielded a main effect of sound frequency (*χ*^*2*^*(12)* = 762.53, *p* < 0.001) and an interaction of group and sound frequency (*χ*^*2*^*(12)* = 26.99, *p* = 0.008), with no main effect of group (*χ*^*2*^*(1)* = 0.7, *p* > 0.05). The linear mixed-effects model on the hardness of sound-induced touch yielded a main effect of sound frequency (*χ*^*2*^*(12)* = 1968.6, *p* < 0.001) and an interaction of group and sound frequency (*χ*^*2*^*(12)* = 194.6, *p* < 0.001), with no main effect of group (*χ*^*2*^*(1)* = 3.11, *p* = 0.078). The linear mixed-effects model on the dryness of sound-induced touch yielded a main effect of sound frequency (*χ*^*2*^*(12)* = 38.4, *p* < 0.001) and no main effect of group (*χ*^*2*^*(1)* = 0.18, *p* > 0.05). The interaction of group and sound frequency approached significance (*χ*^*2*^*(12)* = 20, *p* = 0.067). The linear mixed-effects model on the smoothness of sound-induced touch yielded a main effect of sound frequency (*χ*^*2*^*(12)* = 438.4, *p* < 0.001) and an interaction of group and sound frequency (*χ*^*2*^*(12)* = 73.6, *p* < 0.001), with no main effect of group (*χ*^*2*^*(1)* = 0.002, *p* > 0.05). The linear mixed-effects model on the coldness of sound-induced touch yielded a main effect of sound frequency (*χ*^*2*^*(12)* = 204.1, *p* < 0.001) and an interaction of group and sound frequency (*χ*^*2*^*(12)* = 137.2, *p* < 0.001), with no main effect of group (*χ*^*2*^*(1)* = 0.0, *p* > 0.05). The linear mixed-effects model on the dullness of sound-induced touch yielded a main effect of sound frequency (*χ*^*2*^*(12)* = 3139.3, *p* < 0.001) and an interaction of group and sound frequency (*χ*^*2*^*(12)* = 349.24, *p* < 0.001), with no main effect of group (*χ*^*2*^*(1)* = 1.85, *p* > 0.05). The group by sound frequency interactions for each of the characteristics was due to rating differences for the different sound frequencies in the synesthetes, whereas the controls reported similar ratings across the sound frequencies.

**Figure 3:**
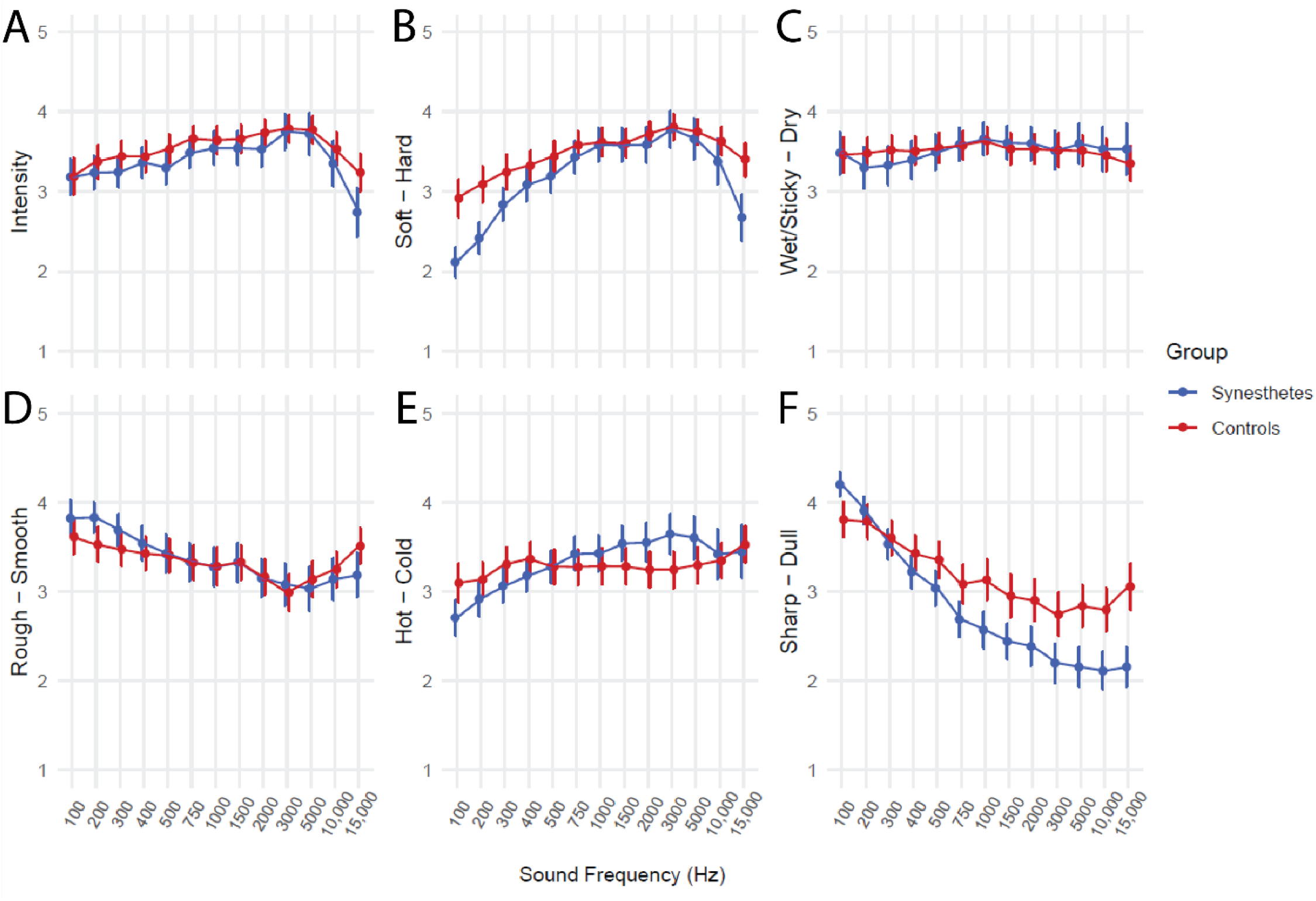
Qualitative ratings of sound-induced touch for synesthetes and controls. Mean intensity (A), hardness (B), dryness (C), smoothness (D), coldness (E) and dullness (F) ratings are represented for each group across sound frequencies. Error bars represent the standard error of the mean.

## Discussion

The primary aim of this study was to characterize and compare the subjective mappings between auditory perception and sound-induced tactile perception in both sound-touch synesthesia and ordinary perception. To do so, we measured the correspondence between sound frequency and the subjective characteristics of induced tactile sensations in synesthetes and control subjects. First, we showed that both synesthetes and controls felt touch when responding to sounds, although synesthetes felt touch more frequently. Second, we showed that both synesthetes and control showed a positive correlation between the location of touch on the body (specifically, height on the longitudinal axis of the body) and sound frequency. However, this correlation was significantly stronger for synesthetes. Third, we showed that the sound-induced tactile sensations felt by synesthetes were more highly concentrated on the body compared to those felt by control subjects. That is, for a given sound frequency, the elicited tactile sensations across multiple presentations of the same sound tended to occur within smaller areas on the body in synesthetes. Finally, we found that sound frequency influenced several qualitative characteristics of touch; specifically with the perceived hardness, roughness, and sharpness of touch in both synesthetes and controls. However, sound frequency did not affect the perceived intensity, temperature, or dryness of touch in either synesthetes or controls. Together, these results suggest that synesthesia is an exaggeration of ordinary perception, since both groups show similar effects, albeit weaker in the control group. Furthermore, the systematicity of the observed behavioral mappings between sound and touch implies the existence of underlying anatomical mappings between somatosensory and auditory areas, and these mappings may be more robust in synesthesia, as suggested by the increased reliability and systematicity of its auditory-tactile correspondences.

This is the first study to date that has systematically characterized sound-touch associations in congenital synesthetes. As such, no established consistency tests exist to formally recognize this type of synesthesia. To do so, we first demonstrated that synesthetes reported feeling touch significantly more frequently in response to sounds compared to controls. We subsequently demonstrated different spatial distributions for the locations of sound-induced tactile sensations in synesthetes and controls. That is, we showed that synesthetes reported feeling sound-induced touch in significantly more concentrated locations on their body compared to controls. This higher consistency in the location of sound-induced touch may be the analog of the high consistency between color and number/letter mappings in grapheme-color synesthesia.

These results demonstrate systematic correspondences between sound frequency and several perceptual dimensions of touch. The correlation between sound frequency and the longitudinal body location of induced feelings of touch in both synesthetes and controls suggest the involvement of somatosensory activations in response to sound. Indeed, the striking pattern of adjacent longitudinal locations for sounds as a function of increasing frequency suggests that sound-induced touch is constrained by the anatomical organization of the somatosensory cortex and, in particular, relies on areas with a somatotopic organization, such as S1 and S2^16–18^. Considering that the auditory cortex also has a topographic organization according to sound frequency (i.e. it has a tonotopic arrangement)^8–10^, it may be hypothesized that functionally organized anatomical pathways between tonotopic and somatotopic representations could underlie the correlations observed between sound and touch in both synesthetes and controls.

In fact, the remarkable correlations observed between sound frequency and the longitudinal location of touch in both groups may rely on the cross-modal pathways that have already been mapped in ordinary perception. Indeed, both behaviorally and anatomically, there is a strong homology between auditory and tactile perception. Since auditory and somatosensory stimuli are pressure-based and their receptors are mechanosensory, they may share similar transduction mechanisms. One study showed that tactile acuity is optimally improved by a concurrent auditory stimulus if it is ipsilateral, and its frequency is congruent with the tactile stimulus^21^ implying the involvement of ipsilateral pathways and homologous mechanisms processing concurrent tactile and auditory information. Consistent with this interpretation, a twin study showed that the same genetic factors contribute to touch sensitivity and hearing^22^.

Anatomically, auditory and somatosensory cortices are adjacent, and imaging studies have shown overlapping responses to sound and touch in perisylvian regions ^23–26^. Cortico-cortical pathways are known to exist between auditory and somatosensory regions. Bidirectional projections between A1 and S1 have been shown in gerbils^27^. In humans, ipsilateral connections between A1 and S1 have also been demonstrated^7^. S2 has also been identified as a site of integration between tactile information and information stemming from other modalities, including vision and audition^28^. Therefore, both behavioral and anatomical evidence demonstrate the existence of interconnectivity between auditory and somatosensory cortices in perception.

One prominent theory of synesthesia ascribes the existence of synesthetic behavioral mappings to the disinhibition of pre-existing neural connections between modalities. In this disinhibited feedback model of synesthesia, the pathways linking the inducing and evoked stimuli, which exist in all people but are normally inhibited, are thought to be disinhibited ^29,30^. In the only neuroimaging studies of sound-touch synesthesia to date, activations in S2^6^, likely resulting from increased connectivity between auditory cortex and somatosensory cortex^7^, were associated with sound-induced tactile sensations. It was hypothesized that a stroke-induced lack of somatosensory thalamic input allowed the unmasking of pre-existing cross-modal connections between adjacent auditory and somatosensory cortices^6^, in line with the disinhibited feedback model. These results have not been corroborated by a study with a larger sample size nor by a study examining congenital synesthetes, whose brains had not undergone post -stroke reorganization.

Together, the present results suggest the existence of functionally specific neural mappings between auditory frequency representations and somatotopic representations. The stronger correlations between sound and touch in synesthetes compared to control subjects, as well as the higher consistency in the location of touch in synesthetes, further suggest that synesthesia could arise through an increased usage of these pre-existing pathways. Future studies should aim to obtain diffusion tensor imaging (DTI) and high-resolution functional magnetic resonance imaging (fMRI) data to test these hypotheses.

An alternative interpretation of these results is that the systematic correspondences between sound and touch are the result of metaphorical and/or semantic mappings that are not perceptual in nature. For example, low versus high pitched sounds may have spatial connotations. Indeed, one study showed that responses to the written word “high” were faster if a high-pitched tone was concurrently played, and similarly with the word “low” and low-pitched sounds^31^. On these grounds, the “SMARC” effect was defined, whereby high-frequency pitches favor up responses and low-frequency pitches favor down responses^32^. This effect could be responsible for the correlation we find between sound frequency and the height of elicited tactile sensations on the body. To test this, a future study could vary the position of the body presented to subjects in each trial, or include different subject positions (e.g., supine vs. upright), to assess whether there are modulations in these correspondences. However, regardless of whether the SMARC effect is responsible for these auditory-somatosensory correspondences, these results are nonetheless strongly suggestive of a perceptual mechanism tying tonotopy with somatotopy. Indeed, whether auditory representations activate somatosensory representations directly, by way of functionally specific anatomical mappings, or via higher-order semantic, spatial, or conceptual representations that activate somatosensory representations indirectly, these results are highly suggestive of a perceptual mapping existing in ordinary and synesthetic perception. In support of this interpretation, one study found that the spontaneous and automatic mappings between sound pitch and the visual features of location, size and spatial frequency in ordinary perception were likely to be perceptual in nature^33^. Furthermore, we also found strong correlations between sound frequency and the perceived roughness, hardness, and sharpness of tactile sensations in both synesthetes and controls. A study of audio-tactile mappings in normal observers found strong associations between the pitch and loudness of sound and specific dimensions of touch, including roughness, sharpness, heaviness and hardness ^34^. Our results in the control participants corroborate these findings, and the presence of these effects in synesthetes further implies that synesthesia relies on the recruitment of the cross-modal pathways that underlie perception in controls. However, the neural mechanisms representing these qualitative dimensions of touch perception are poorly understood, and more research will be necessary to characterize them.

In conclusion, this study is the first to demonstrate systematic relationships between auditory and tactile perception in both congenital synesthesia and ordinary perception. This significant behavioral mapping suggests underlying neural mappings between audition and touch, which may cause the sound-induced somatosensory activations that mediate the tactile sensations felt by synesthetes. These findings emphasize the importance of studying the neural mechanisms of synesthesia and their potential in improving our understanding of normal brain function.

## Materials and Methods

This research was approved by the Institutional Review Board of the City University of New York.

### Subjects

Sixty-two synesthetes were recruited from synesthesia databases and online synesthesia forums to complete a screening test (described below) for sound -touch synesthesia. Of these participants, fifty-two were invited to participate in the experiment. Twenty-two did not respond to the invitation, seven indicated that the sounds used in the screener were too uncomfortable or painful, and four had logistical problems that prevented their participation. As a result, nineteen subjects (14 females, 4 males, 1 other; mean age 36.1, range: 18 – 67; 14 right-handed, 1 left-handed, 4 ambidextrous) participated after informed consent. Twenty-three control subjects were recruited online from Amazon Mechanical Turk (6 females, 17 males; mean age 36.4; range: 20 – 57; 20 right-handed, 2 left-handed, 1 ambidextrous) and participated after informed consent.

For the synesthesia group’s screening test, which was conducted upon initial recruitment, each subject was sent a link to a web-based screener that played a total of forty sounds in random order, such that each of thirteen different sounds was played at least three times. For each sound, subjects were asked “Did you feel this sound?” and responded by selecting one of the following options: “No”, “Very Weakly”, “Weakly”, “Moderately”, “Strongly” or “Very Strongly”. They were then asked to indicate where they felt the sensation on their body by selecting one or more of the following options: “Left Leg”, “Left Arm”, “Left Face”, “Right Leg”, “Right Arm”, “Right Face”. After the screening was completed, subjects who selected the same intensity and/or location of the tactile sensation for three iterations of the same sound, for at least six sounds, were invited to take part in the main experiment.

### Stimuli

The stimuli used for the screening test and the main experiment were identical. Thirteen one-second-long sounds were produced using Matlab (Mathworks, Natick, MA) at frequencies of 100 Hz, 200 Hz, 300 Hz, 400 Hz, 500 Hz, 750 Hz, 1000 Hz, 1500 Hz, 2000 Hz, 3000 Hz, 5000 Hz, 10 000 Hz, and 15 000 Hz. All sounds were played continuously on loop and were corrected for loudness^35^.

### Experimental Paradigm

In each trial, one of the thirteen sounds played automatically and repeatedly until the subject indicated whether they could feel the sound as a tactile sensation or not (see Figure 1A). If they did, the sound repeated while they answered the following seven questions: first, they indicated by a click where on an image of a gender-neutral body and face they had felt the sound. Subjects were asked to indicate the location with the strongest tactile sensation, even if the sensation was large or diffuse. Second, subjects indicated the intensity of the sensation on a scale of 1 to 5. Finally, subjects answered five questions about the qualitative aspects of the sensation, by rating the following measures on a scale of 1 to 5: soft to hard, wet/sticky to dry, rough to smooth, hot to cold, and sharp to dull. For each of the seven questions, subjects were given the option to select “not applicable”.

Each of the 13 sounds was repeated 20 times, with the exception of the 1500 Hz sound, which was repeated only 19 times due to an error in the software preparation. There was a total of 259 trials, with the order of sounds randomized within 19 blocks of 13 trials and one block of 12 trials. All stimuli and response collection were hosted and controlled using custom software hosted by Qualtrics (Provo, UT).

### Data Analysis

The XY pixel coordinates selected by subjects to indicate the location of tactile sensations on the body diagram were first transformed such that coordinates selected on the back-facing body or on the two face diagrams were mapped to equivalent locations on the front-facing body. These transformations were carried out because our hypotheses pertained only to the lateral (left-right) and longitudinal (upper-lower) locations of tactile sensations on the body and did not predict any relationship between sound frequency and the ventral-dorsal axis. Coordinates selected in the white space around the body or face diagrams were excluded from further analysis.

To characterize differences in sound-induced tactile sensations between the synesthete and control groups, we first measured the rate of sound-induced touch sensations in both groups. We used a binomial generalized linear mixed effects model (GLMM) to assess the effect of group and sound frequency on the rate of tactile sensations.

We then measured the spatial distribution on the body of sound-induced tactile sensations in both groups. To do so, we first aimed to assess whether the location of touch on the body was influenced by sound frequency. To this end, we computed the correlation between sound frequency and the location of touch on the lateral body axis (X coordinates) and between sound frequency and the location of touch on the longitudinal body axis (Y coordinates), in both groups. We used Fisher’s z-transformation to obtain normalized correlation coefficients between sound and location, then used a one-sample t-test against zero to assess whether correlations between sound and location were significant within a group (synesthetes and controls) and along a body axis (lateral, X and longitudinal, Y). Finally, we used two-sample t-tests to assess whether correlations between sound and location were different across groups (synesthesia versus control group, for X-axis and for Y-axis).

In a second analysis, we aimed to assess whether different sounds were associated with touch in more concentrated areas of the body across groups. To do so, we converted the locations of sound-induced touch into polar coordinates and computed the mean resultant length, a measure of the spatial concentration of the selected locations on the body in response to each sound frequency. A linear mixed effects model (LMM) was computed with subjects as a random effect, to assess the effects of group and sound frequency on the mean resultant length.

Finally, we used linear mixed effects models to investigate the effects of group and sound frequency on the subjective intensity, hardness, dryness, smoothness, coldness and dullness of sound-evoked tactile sensations. All p-values were false discovery rate (FDR)-corrected for multiple comparisons.

## Acknowledgments

This work was supported by the Doctoral Student Research Grant of the Graduate Center of the City University of New York to LK and by NSF BCS Awards #1358893 and #1755477 and NSF EFRI Award #1137172 to TR.

